# Teaching data science fundamentals through realistic synthetic clinical cardiovascular data

**DOI:** 10.1101/232611

**Authors:** Ted Laderas, Nicole Vasilevsky, Bjorn Pederson, Melissa Haendel, Shannon McWeeney, David A. Dorr

## Abstract

**Objective:** Our goal was to create a synthetic dataset and curricular materials to assist in teaching fundamentals of translational data science.

**Materials and Methods:** A literature review was conducted to extract current cardiovascular risk score logic, data elements, and population characteristics. Then, clinical data elements in the models were pulled from clinical data and transformed to the Observational Medical Outcomes Partnership (OMOP) common data model; genetic data elements were added based on population rates. A hybrid Bayesian network was used to create synthetic data from the logical elements of the risk scores and the underlying population frequencies of the clinical data.

**Results:** A synthetic dataset of 446,000 patients was created. A two-day curriculum was created based on this synthetic data with exploratory data analysis and machine learning components. The curriculum was offered on two separate occasions; the two groups of learners were given the curriculum and data, and results were tallied, summarized, and compared. Students’ ability to complete the challenge was mixed; more experienced students achieved a range of 70%-85% in balanced accuracy, but many others did not perform better than the baseline model.

**Discussion:** Overall, students enjoyed the course and dataset, but some struggled to consistently apply machine learning techniques. The curriculum, data set, techniques for generation, and results are available for others to use for their own training.

**Conclusion:** A realistic synthetic data with clinical and genetic components helps students learn issues in cardiovascular risk scoring, practice data science skills, and compete in a challenge to improve identification of risk.

## BACKGROUND AND SIGNIFICANCE

The promise of data and information - and their related systems - to improve health and well-being has grown in recent years, but the ability to use analytics effectively to harness this potential has lagged behind the promise. For instance, heart disease remains the top cause of death in the United States, [1] accounting for over 30% of yearly deaths, yet optimal prediction of cardiovascular risk and addressing risk factors can lower the odds of a heart attack by 80%, of a stroke by 69%, and overall mortality by 45%. [2] Segmenting population by risk, however, has been slow; risk prediction scores such as Framingham, the Atherosclerotic Cardiovascular Disease (ASCVD) risk score, and the Mesa score all predict well at the population level, but miss crucial subpopulations such as younger persons, those of minority ethnic or racial backgrounds, or those with genetic risks. [3]

For experienced data scientists, there is poor access to real clinical data that can be used to train new classifiers or to develop new algorithms. Synthetic data generation approaches attempt to address these issues by using real data as a basis for generating and modeling synthetic approaches. [4–6] Additionally, despite an unprecedented level of investment in prediction capability for cardiovascular disease fe.g, [7]), researchers often lack fundamental skills in analytics, including exploratory data analysis (EDA), assessing data quality and fitness for the problem at hand, data curation and integration; these gaps lead to significant issues in the validity and reproducibility of research results. [8,9]

As part of the NIH Big Data to Knowledge (BD2K) initiative, we have developed a synthetic dataset and coursework to teach students the difficulties of working with both clinical and genetic data for prediction. These difficulties include inadequate phenotype mapping, the impact of comorbidities, and low disease prevalence in patient cohorts.

In light of these difficulties, a growing concern is that students do not spend enough time doing exploratory data analysis (EDA) in order to understand how these difficulties can impact their analytical models. Successful applications of machine learning approaches to domains such as clinical and genetic data require students to gauge the difficulty of the problem, and understand whether assumptions of the algorithms (such as a balanced cohort for Random Forests) used are met for a particular data set. In short, critical thinking, domain knowledge, and curiosity are required for effective application of machine learning.

In order to facilitate development of these skills, we have developed a synthetic dataset and curricular materials for a 2 day workshop modeled on a real patient cohort that includes clinical and genetic covariates. This dataset includes realistic dependencies between variables (such as high BMI patients having a higher prevalence of Type 2 diabetes) as well as low disease prevalence for certain cohorts (such as those younger than 40). This dataset can be used for multiple purposes: 1) for teaching math, statistics, and computer science students the challenges of using real world data, 2) teaching machine learning techniques to biologists and clinicians, and 3) assessing the performance of machine learning techniques on a tunable dataset.

## OBJECTIVE: TEACHING ROBUST RISK PREDICTION

Our goal for this effort was to integrate our learning objectives with a realistic synthetic dataset. We chose cardiovascular risk prediction as a problem to model for multiple reasons: 1) high prevalence in the populations of interest, 2) identification of possible genetic factors involved, and 3) the difficulty of predicting risk in cohorts. Overall, most risk calculators do not cover the younger population (20-40 year olds), and we believe that teaching the students the difficulties of predicting risk in low prevalence populations would highlight many of the issues with modeling clinical data. Our goal was to create a dataset that was feasible for novice students to learn on, but realistic enough for students to encounter difficulties with. To this end, we needed to simplify the structure and dependencies in the data.

### Learning Objectives

There were two learning objectives to our workshop: 1) Assess overall risk in the synthetic data and identify a cohort in which to predict CVD risk for, and 2) given this cohort, select appropriate covariates to predict risk using machine learning techniques. The use of active learning methods in our workshop addresses key areas identified for statistics education by both Garfield and Kaplan, such as construction and ownership of learning, allowing students to become aware of their own mistakes and confront errors in their reasoning. [10]

Objective 1 was implemented via a R/Shiny interactive dashboard (see Figure 1) to assess covariates, their interactions, and their predictive value in predicting CVD. Shiny is a lightweight interactive visualization framework written in R and was used to conduct Exploratory Data Analysis of our synthetic cohort. The Shiny interface gives students a high level view of the data, allowing them to identify a cohort of interest and assess associations between variables.

**Figure 1.**
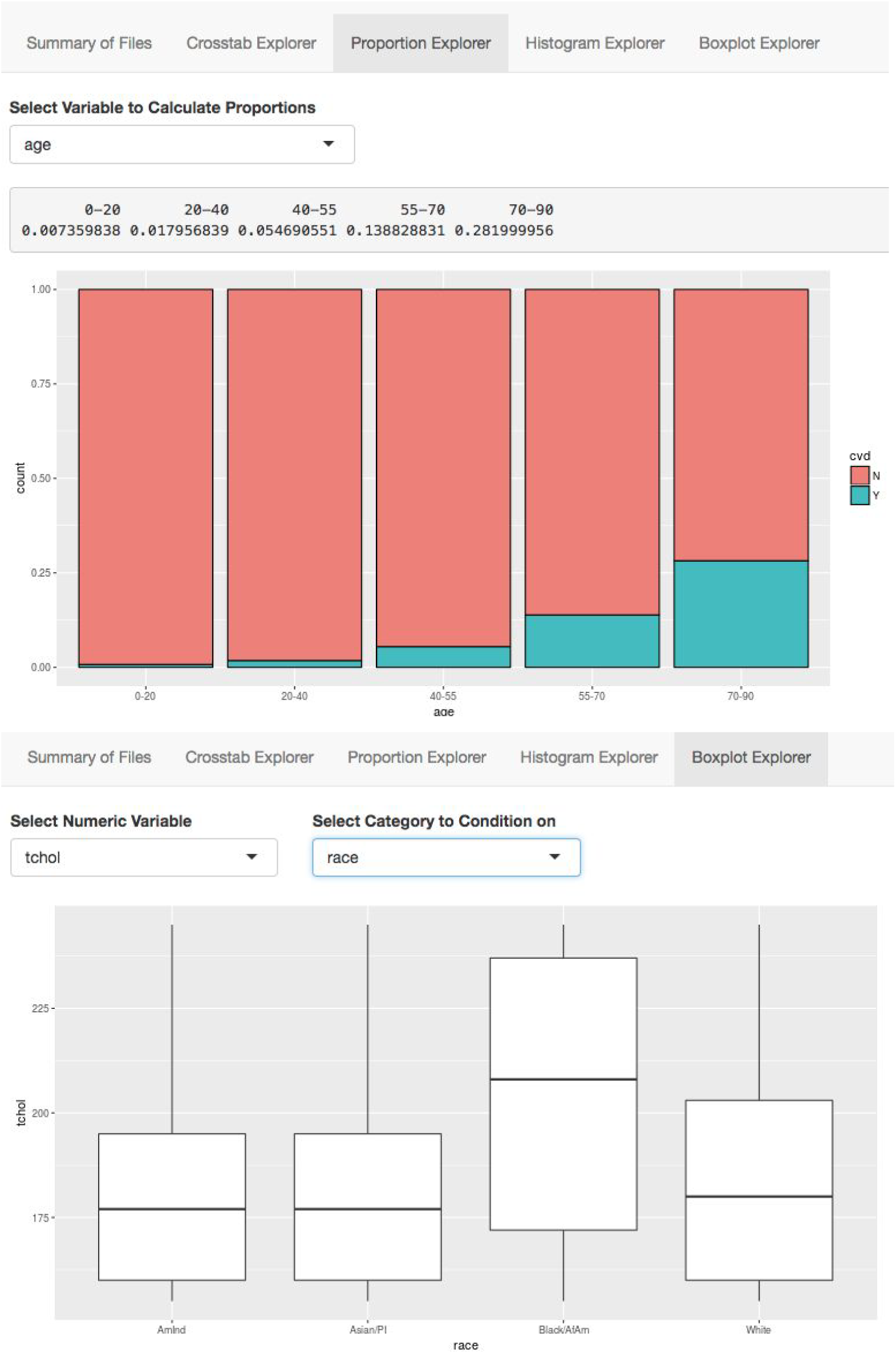
Shiny Visualization Dashboard for Exploratory Data Analysis. a) Proportion explorer allows for visual exploration of covariates (here, age category) with outcome (CVD). b) Boxplot Explorer for exploration of continuous covariates (here total cholesterol) conditioned on categorical covariates (here, race) in order to assess interactions between covariates.

Objective 2 was implemented as a simplified programming exercise in R using three different modeling techniques (logistic regression, linear discriminant analysis, and classification and decision trees). In order to facilitate discussion, we used an online scoreboard so students could compare their cohorts and the performance of their models on those cohorts. We followed up the modeling results with a discussion of their findings and the true structure of the data.

We targeted the workshop to students and staff at our neighboring university (Portland State University), who had basic familiarity with R and statistics. The majority of the audience who signed up were graduate students (86.4%), but also included some undergraduate students (9.1%) and one staff member (4.5%). We also assessed whether participants had knowledge of statistical techniques such as t-tests, ANOVA/linear modeling, and multivariate regression.

Additionally, we used a reduced set of the covariates and data as a modeling exercise for students in our data analytics course, whose backgrounds ranged from clinical (including cardiology and emergency medicine) to those with computational biology backgrounds. We expected overall better metrics from this second group, due to their domain knowledge and greater familiarity with data analysis.

## MATERIALS AND METHODS

### Mapping clinical cohort to a common data model

We used clinical data from a OHSU cohort to seed the clinical data. Specifically, we extracted data from patients who had at least yearly visits with primary care or medical subspecialities in ambulatory clinics over a 2 year period. We chose concepts to extract based on a review of the data needed for 3 common risk models (ASCVD, Mesa, and the updated Framingham models). Key concepts from demographics (age, gender, race, ethnicity), diagnoses, medications (antihypertensives), vital signs (Body Mass Index, or BMI; Systolic and diastolic blood pressure), and laboratory values (Total, High density, and low density cholesterol) were extracted and, where necessary, coded into standard terminologies and groups that appeared to match the risk score definitions. We then transformed the source clinical data into the Observational Medical Outcomes Partnership (OMOP) common data model maintained by the Observational Health Data Sciences and Informatics (OHDSI) collaborative [11] to our cohort in order to assess our cohort data quality and to utilize pre-made queries to pull covariates appropriate to our synthetic data. To assess data quality, we ran the `Automated Characterization of Health Information at Large-scale Longitudinal Evidence Systems (ACHILLES)` tool maintained by OHDSI using R; we addressed any significant findings from ACHILLES. [12] Then, we further transformed the data to simplify the structure for teaching.

### Using a hybrid Bayesian network approach to generate consistent clinical and genetic data

A Bayesian Network (Figure 2) was used to encode the dependencies between variables in the dataset, such as patients who do not have hypertension (Hypertension = N) should not receive hypertension treatment (Treatment = N). Where possible, we have utilized frequency tables from our patient cohort to generate the conditional probability tables (CPTs) for variables. When this information is not available, incidence probabilities (such as smoking given age, probability of receiving hypertension treatment) were derived from relevant CDC and NHLBI studies. [13] Our generalized workflow is available at the cvdRiskData repository (http://github.com/laderast/cvdRiskData).

**Figure 2.**
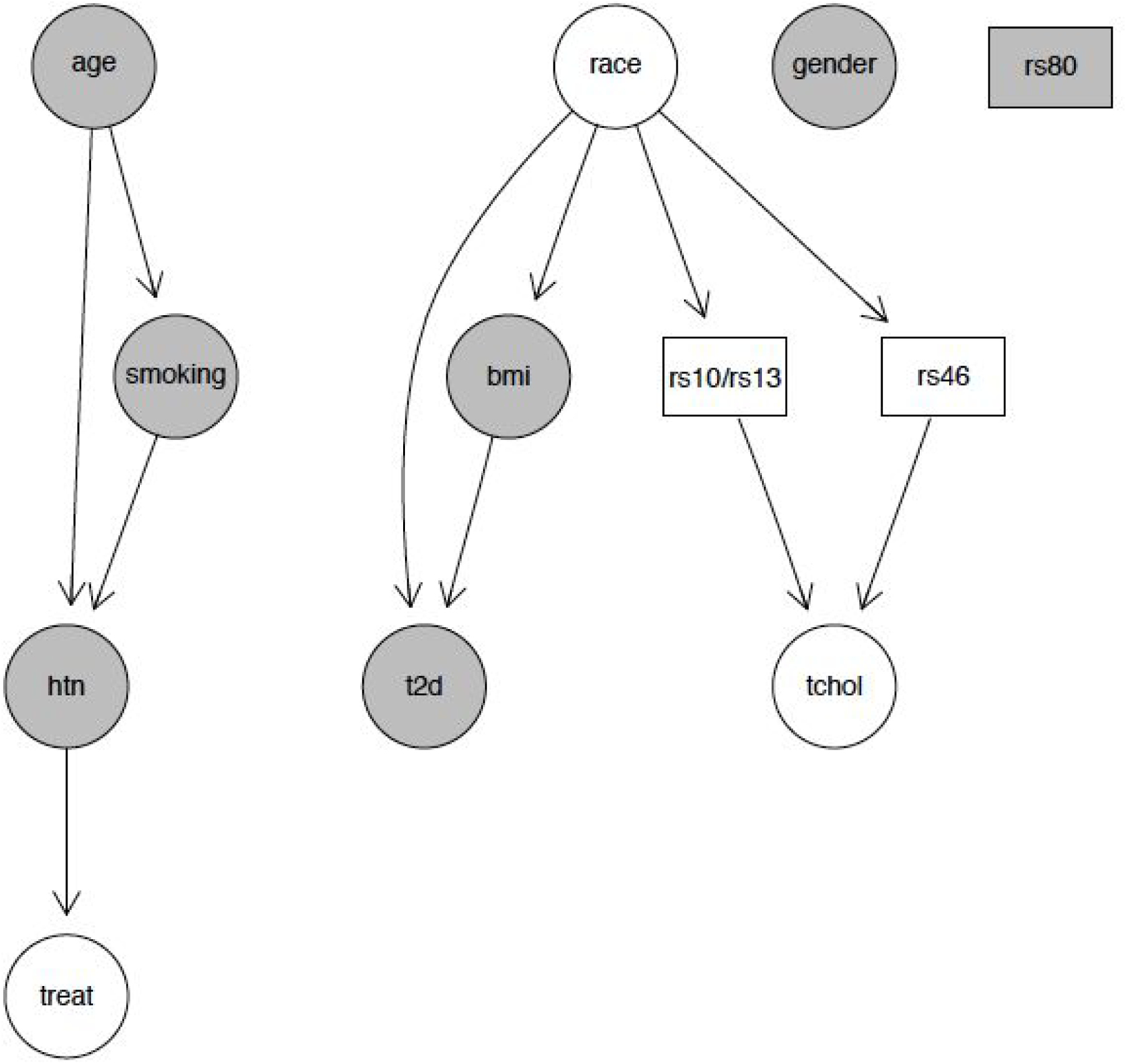
Bayesian Network Structure of dataset. Arrows indicate dependencies, modeled with conditional probability tables. Nodes in grey directly influence overall risk in dataset (calculated as variation on Framingham risk score), whereas white nodes (`nace, treat` (hypertension treatment), `tchol` (total cholesterol)) have indirect effects on risk. Categorical data is transformed to continuous data via sampling within a categorical interval. Boxes are genetic component of data, calculated but only supplied for 10% of original dataset. Presence of rs80 homozygous variant mediates risk in entire patient cohort, increasing risk four-fold for a patient.

In order to simplify the problem, we limited our variables to have no more than five states each. We also limited the number of dependencies to any variable in the model to be 2 or less, a common assumption when building Bayesian Networks. These restrictions resulted in a Bayesian Network encoding three sets of dependent variables (*Age-Hypertension-Smoking-Treatment, Race-Genotype-Bmi-TChol-T2D, Gender, Figure 2*). These three sets of variables were considered conditionally independent of each other. The Bayesian Network was implemented using the *gRain* package in R. *gRain*’s `simulate`() function allows for the generation of arbitrarily large datasets from a specified Bayesian network.

In our dataset, we needed to simplify the overall dependencies in the data. Overall, CVD risk is mediated through the five variables used in the Framingham Risk score (age, systolic blood pressure, hypertension treatment, BMI, Type 2 Diabetes status). [14] Other available covariates (race, gender, smoking status, genotype) indirectly influence these variables. For example, smoking status affects hypertension, but is not directly linked to CVD risk (Figure 2).

Implementing the above Bayesian networks resulted in marginal distributions that agreed with our data and US-wide distributions (see validation section below). One difficulty we encountered was that attempting to integrate the five risk factors to calculate the probability of CVD with our data would have required a large CPT, requiring us to estimate 1024 conditional probabilities. Instead of coding a large CPT to integrate all five risk factors, we first converted categorical data to continuous variables where necessary, and then implemented the Framingham risk score to assess risk in the appropriate variables. For each synthetic patient, we calculated risk and used this risk as a probability that patient had a 10 year CVD risk. We then assessed the overall CVD prevalence in our synthetic population, which was high compared to our real data. In order to address this, we then downsampled the number of CVD cases in our dataset to adjust the prevalence of our dataset. Overall, our data has a higher prevalence (8.7%) compared to our observed data.

### Integration of Genetic Covariates

We integrated the genetic covariates as an aggregate genotype of 4 SNP variants (Figure 2). Because of the limited workshop time, we chose just 4 SNPs to teach students about the basics of genetic structure and inheritance. Two of these SNP variants (rs10757278 and rs1333049) were chosen to co-occur (due to linkage disequilibrium), providing the same information. We assigned three SNPs (rs10757278, rs1333049 and rs4665058) to be associated with elevated total cholesterol in our dataset (by providing the appropriate CPT), and one SNP (rs8055236) directly influences overall risk (homozygous variant = 4 x Risk). Individual SNP genotypes were limited to either homozygous wild-type or variant. To simplify the problem, we limited the number of observed aggregate genotypes to six overall. The frequency of each SNP was dependent on race (derived from SNPedia [15]), and we calculated the aggregate genotype frequency for each race by multiplying the individual SNP probabilities for each race. This resulted in different aggregate genotype frequencies across the cohort. Essentially, for three of these SNPs, genetic contribution to the risk score was mediated through other variables (total cholesterol and race), with only one SNP directly mediating risk in the total population.

We limited supplying these genetic covariates to a smaller cohort, about 10% of the original population. We did this to simulate the likelihood of a patient receiving a genetic screen. However, the full dataset includes risk that is mediated by these genetic covariates. Because of these missing covariates, building a predictor to predict CVD risk perfectly is not possible for the larger dataset.

### Validation of synthetic dataset

We used an iterative process for validating the realism of the clinical dataset. Once the data was generated, we examined marginal probabilities to ensure they fit our expectations, and visualized the data using our dashboard (see below) in order to assess the strength of the associations in our data. A critical part of the validation process was achieved by identifying key questions that students might have of the data, such as “Is there an association between race and BMI?”, and assessing whether these questions had realistic associations in our dataset.

### Day 1 Workshop: Interactive visualization for exploratory data analysis

After a brief overview of cardiovascular risk prediction scores, we had the students explore the dataset to identify a smaller cohort of interest. In order to accomplish this, we implemented a dashboard for examining the data using R and Shiny, an interactive visualization framework. The dashboard enables students to ask questions of association using a number of helpful tools (such as data summaries and crosstables) and visualizations (such as histograms and boxplots). For example, the Proportion Explorer allows students to ask questions of association of our outcome variable (CVD risk) with categorical variables, such as (ageCategory, hypertension, or race) (Figure 1a). Boxplots allow students to stratify the continuous data (`sbp, numAge`) by categorical variables (Figure 1b). A version of the EDA dashboard (including the synthetic dataset) is available here: https://tladeras.shinyapps.io/cvdnight1/.

### Day 2: Assessment of Machine Learning on a selected cohort

We gave the students the options of using 3 machine learning methods (logistic regression, linear discriminant analysis, and classification and regression trees (CART)) to predict CVD risk in their selected cohort (examples include patients over 55, women over 60, and patients under 40). Students were supplied with Rmarkdown documents with R code to modify for this purpose. [16] We required students to justify why they included specific covariates in the model. One advanced student used a different machine learning tool, SuperLearner, for their modeling of the cohort. [17] We used a scoreboard (implemented as a Google Sheet) to compare predictions by cohort prevalence using three different metrics: *sensitivity, positive predictive value* (PPV), and *balanced accuracy*. We chose these metrics because they favor the low prevalence positive case (CVD=1), whereas metrics like *accuracy* favor the high prevalence negative cases (CVD=0) (Figure 3).

**Figure 3.**
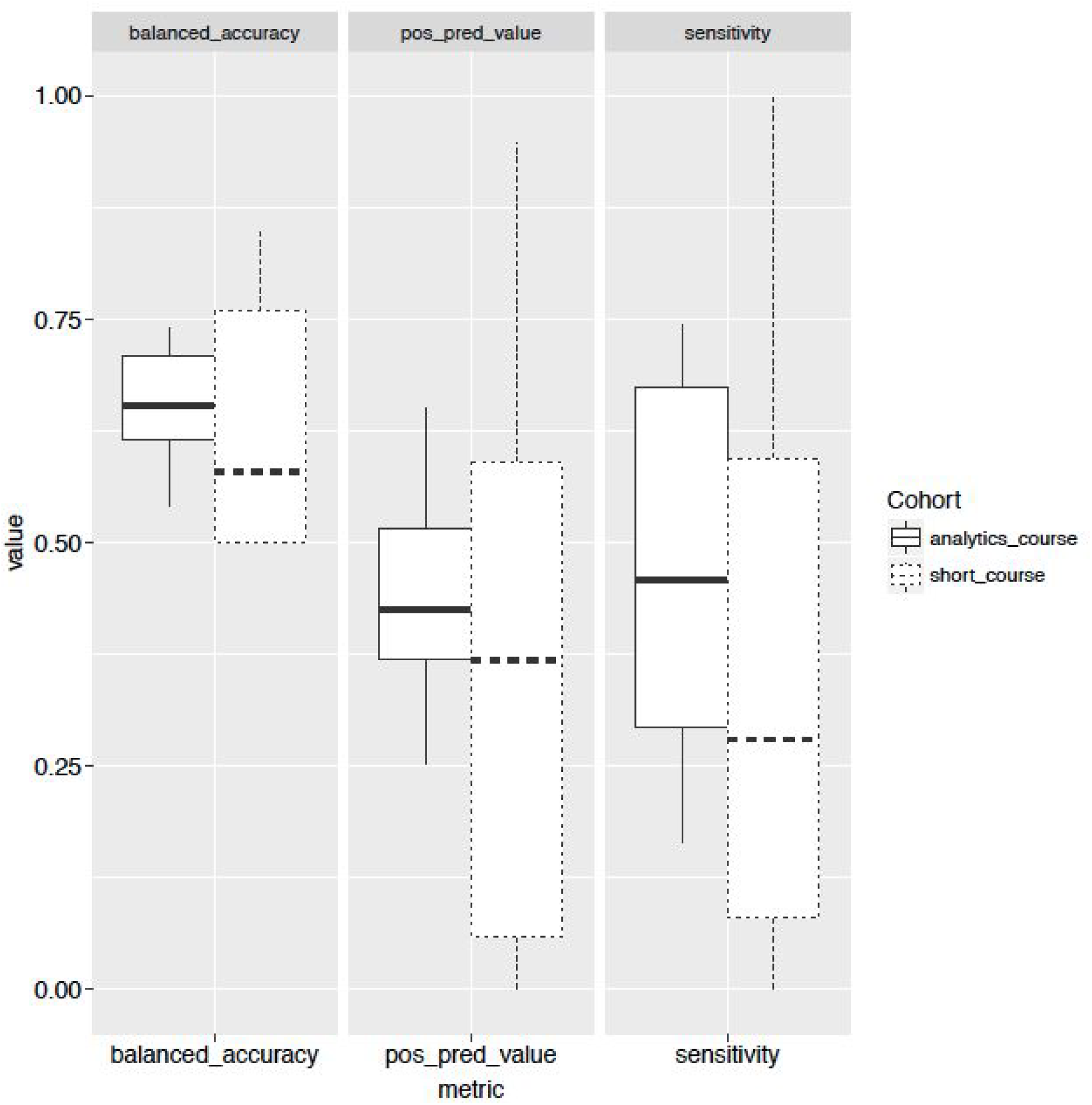
Metrics for Short Course/Analytics Course. Comparison of metrics of Short Course and Analytics Course students. Shown are 21 logistic regression models for the analytics course and 11 models built for the short course (with different probability thresholds) of students in Analytics class when asked to predict CVD risk with subset of dataset (dataset did not include genetic covariates).

## RESULTS

### Demographics of our patient cohort and synthetic data

Table 1 shows the descriptive statistics of the populations used to estimate each risk score, compared with our clinical dataset, and the final synthetic cohort. The initial clinical data had significantly lower frequencies of the key outcome (heart attack or stroke), had a larger proportion of caucasian patients, and lower rates of reported Hispanic ethnicities. These matched local clinical populations. The synthetic data had similar proportions to the clinical data with variations < 10% of means.

**Table 1.**
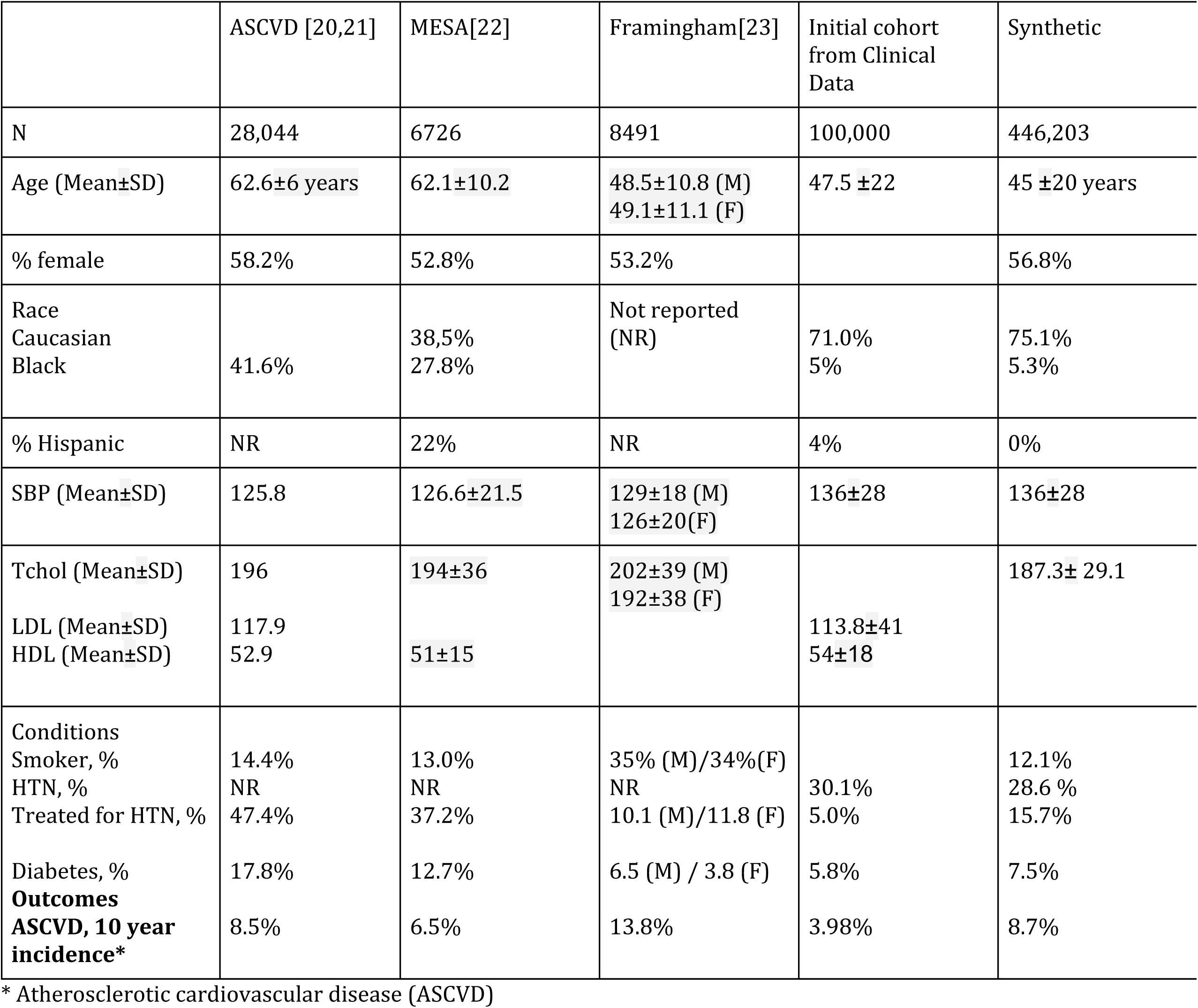
Descriptive statistics of key cardiovascular risk prediction studies, the original clinical data, and the synthetic data set

### Examination of variable dependencies in data

We show the clinical covariates with the dashboard (https://tladeras.shinyapps.io/cvdnight1/) so that interactions between variables can be examined. As expected, hypertension treatment (treat) is not done for those who do not have hypertension. As age category increases, there is a higher prevalence of CVD disease. In high risk CVD patients, there is a higher mean BMI. In building the dataset, examining the distribution of outcomes among our categorical variables was a valuable sanity check.

### Modeling results from both student cohorts

We show results from both the short course and the analytics course students (Figure 3). Overall, for the short course students, which included the cohort selection step, positive predictive value of their models was correlated with prevalence of CVD in their selected cohorts. A few of the models built by the students in this cohort did not converge (such as Cohort: HTN and covariates: smoking, or Cohort: Age = 20-40 years), indicating that the student may have lacked understanding of the inherent issues in the modeling. Two students did tackle the genetic covariates and higher sensitivity and balanced accuracy than the others, which was as expected.

Analytics course students, who received more training in logistic regression and clinical backgrounds, had higher overall mean sensitivity, positive predictive value, and balanced accuracy than the short course students, indicating that more training and domain knowledge led to better overall models and results.

## DISCUSSION

### Mapping Clinical Covariates from OHDSI

Mapping clinical covariates, such as estimating number of hypertension cases, number of Type 2 Diabetes cases, was challenging even given value sets for these diseases. Much of this difficulty may have arisen from the mapping process into the OHDSI model. The original clinical data had not previously been mapped into a common data model, and the ability to identify all instances of the important concepts from the data may have been incomplete, either through issues mapping or through data incompleteness. These difficulties in mapping into the common data model resulted in underestimates for many of our marginal probabilities; however, they do provide a realistic problem for learners to address, as incompleteness and potential inaccuracies in data quality have limited generalizability of many models.

### Lessons learned in synthesizing the data

Implementing the Bayesian Network required multiple levels of tuning and adjustment. One complication was that we were unable to estimate many of the CPTs required by our network from our clinical dataset. In our case, we used estimates of prevalence for the population of the United States. Additionally, we simplified our network considerably to three separate networks to produce our data. Many of the interaction probabilities, such as Age/Hypertension, and Race/BMI, had CPTs that had to be carefully tuned by hand. Ideally, we would have estimated these parameters from the original clinical dataset. [18]

Since producing a CPT integrating the five types of influential covariates was difficult, we opted for combining the covariates using one of the actual risk prediction equations (Framingham [14]). Using a logistic model to combine covariates allowed us to tune the difficulty of the model by adjusting predicted probabilities. Additionally, it allowed us to integrate the effects of one of the SNPs to increase overall risk in a realistic fashion. We believe that this hybrid approach was appropriate for the problem.

The structure of the data included a genetic component, which influenced overall risk. However, this component was not included as a covariate in the larger dataset. Thus, building a machine learning or statistical model that perfectly predicts the data is not possible with the larger dataset. We believe that this missing explanatory variable actually makes the exercise more realistic, as many unmeasured covariates (such as social determinants of health) may influence overall risk.

### Workshop and Modeling Results

Based on student feedback, we believe that the students who participated in the two-day workshop found it informative and challenging, which is also indicated with the overall scores for their model. The distribution of our modeling results indicates that the dataset is a challenging, but not too difficult problem, especially for students who understand the importance of disease prevalence in the data.

Students in the short course, who had mainly mathematics and computer science backgrounds, struggled somewhat with understanding the importance of particular covariates in prediction. We plan to address this with an exercise in understanding the importance of these covariates in CVD. Some students did build models that did not converge, showing a lack of understanding of the inherent issues with the dataset. However, we believe that these models are teachable moments, allowing us to assess their understanding of the issues and provide more information to the students at the time of the workshop. Overall, the analytics students, who had more clinical or machine learning backgrounds, selected more highly predictive models than the short course students. However, one student in the short course had very high scores for PPV and Sensitivity for their models. Following up with this student, we found that they had used an ensemble approach, indicating that he was more experienced in the machine learning component.

### Other uses for the dataset

As shown in Table 1, the dataset has realistic distributions over several key variables, and contains realistic outcome assessments for subpopulations of patients defined in the hybrid Bayesian networks. The dataset may be used freely for validating clinical queries, assessing data quality, benchmarking of new machine learning algorithms, and assessing risk of key outcomes. We believe that such synthetic datasets that are free of proprietary EHR systems will prove key in overcoming obstacles to implementing machine learning algorithms in clinical data. In contrast to larger synthetic generation efforts such as SYNTHEA, we do not attempt to model longitudinal data nor the interaction of multiple co-morbidities. [4] Limiting our scope to CVD has helped us make this dataset as realistic as possible.

### Limitations

Our results are limited to the student cohort at Portland State University and informatics students at OHSU, most of which had a fairly strong mathematics or clinical background. We recommend that students have a basic understanding of statistical concepts before attending the workshop. Additionally

### Future Directions

We believe that the data generation framework is highly generalizable to teaching other important data science/analysis skills, such as mapping phenotypes using ontologies, data cleaning, and meta-analysis. Biases can be introduced into the data to represent issues such as batch/site effects, and other clinical data quality issues. [19] We plan on developing more workshops leveraging this approach and making the course material available as additional open educational resources (OERs).

### Data, generation script, and course material availability

The current version of the synthetic dataset is available as an R package called `cvdRiskData` on GitHub (http://github.com/laderast/cvdRiskData). This package also includes the script, Bayesian network, and CPTs used to generate the dataset. Our course materials for teaching the workshop as well as the dataset simulation script are also available (http://github.com/laderast/cvdNight1 and http://github.com/laderast/cvdNight2).

## CONCLUSION

We believe that training students in exploratory data analysis using realistic clinical data is a gap in most current data science curricula. Without such training, it is difficult to understand the issues in modeling real-world data. In order to address this gap, we have created a synthetic dataset modeled on a real patient cohort that presents students challenges in EDA and modeling. This dataset is an opportunity for active learning of real issues in the data, especially in understanding why it is difficult to predict cardiovascular risk in low prevalence populations, such as 20-40 year olds. Our dataset has further utility as a test dataset for machine learning methods on selected cohorts. We have shared our material as Open Educational Resources (OERs), and welcome further refinements to the dataset generation from the informatics and data science community.

## Acknowledgements

We would like to thank Christopher Chute and Harold Lehmann for their discussion and ideas around synthesizing data using Bayesian Networks. This research was funded with a BD2K Training grant: 1R25EB020379-01.

## References

1 Health, United States, 2016. US Department of Health and Services, Centers for Disease Control and Prevention, National Center for Health and Statistics. 2016. https://www.cdc.gov/nchs/data/hus/hus16.pdf

2 Benjamin EJ, Blaha MJ, Chiuve SE, et al. Heart Disease and Stroke Statistics—2017 Update: A Report From the American Heart Association. Circulation 2017;:CIR.0000000000000485.

3 Puckelwartz MJ. The Missing LINC for Genetic Cardiovascular Disease? Circ Cardiovasc Genet 2017;10. doi: 10.1161/CIRCGENETICS.117.001793

4 Walonoski J, Kramer M, Nichols J, et al. Synthea: An approach, method, and software mechanism for generating synthetic patients and the synthetic electronic health care record. J Am Med Inform Assoc Published Online First: 30 August 2017. doi: 10.1093/jamia/ocx079

5 Buczak AL, Babin S, Moniz L. Data-driven approach for creating synthetic electronic medical records. BMC Med Inform Decis Mak 2010;10:59.

6 Choi E, Biswal S, Malin B, et al. Generating Multi-label Discrete Electronic Health Records using Generative Adversarial Networks. arXiv preprint arXiv:1703 06490 Published Online First: 2017.https://arxiv.org/abs/1703.06490

7 One Brave Idea. http://www.onebraveidea.com/ (accessed 13 Nov 2017).

8 Safran C, Shabot MM, Munger BS, et al. Program requirements for fellowship education in the subspecialty of clinical informatics. J Am Med Inform Assoc 2009;16:158–66.

9 Berner ES, Dorsey AD, Garrie RL, et al. Assessment-based health informatics curriculum improvement. J Am Med Inform Assoc 2016;23:813–8.

10 Garfield J, Ben-Zvi D. How Students Learn Statistics Revisited: A Current Review of Research on Teaching and Learning Statistics. Int Stat Rev 2007;75:372–96.

11 Huser V, DeFalco FJ, Schuemie M, et al. Multisite Evaluation of a Data Quality Tool for Patient-Level Clinical Data Sets. EGEMS (Wash DC) 2016;4:1239.

12 ACHILLES for data characterization - OHDSI. https://www.ohdsi.org/analytic-tools/achilles-for-data-characterization/ (accessed 13 Nov 2017).

13 CDC - DHDSP - Heart Disease Facts and Statistics. https://www.cdc.gov/heartdisease/statistics.htm (accessed 14 Aug 2017).

14 D’Agostino RB, Russell MW, Huse DM, et al. Primary and subsequent coronary risk appraisal: new results from the Framingham study. Am Heart J 2000;139:272–81.

15 Cariaso M, Lennon G. SNPedia: a wiki supporting personal genome annotation, interpretation and analysis. Nucleic Acids Res 2012;40: D1308–12.

16 Allaire J, Cheng J, Xie Y, et al. rmarkdown: Dynamic Documents for R, 2016. *R package version 0 9* 2016;6.

17 Polley EC, Van der Laan MJ. SuperLearner: Super Learner prediction, Package Version 2.0-4. Vienna: R Foundation for Statistical Computing 2011.

18 Larsson SC, Burgess S, Michaëlsson K. Association of Genetic Variants Related to Serum Calcium Levels With Coronary Artery Disease and Myocardial Infarction.JAMA 2017;318: 371–80.

19 Weiskopf NG, Weng C. Methods and dimensions of electronic health record data quality assessment: enabling reuse for clinical research. J Am Med Inform Assoc 2013;20:144–51.

20 Goff DC, Lloyd-Jones DM, Bennett G, et al. 2013 ACC/AHA guideline on the assessment of cardiovascular risk. Circulation 2013;:01 – cir.

21 Muntner P, Colantonio LD, Cushman M, et al. Validation of the atherosclerotic cardiovascular disease Pooled Cohort risk equations. JAMA 2014;311: 1406–15.

22 McClelland RL, Jorgensen NW, Budoff M, et al. 10-year coronary heart disease risk prediction using coronary artery calcium and traditional risk factors: derivation in the MESA (Multi-Ethnic Study of Atherosclerosis) with validation in the HNR (Heinz Nixdorf Recall) study and the DHS (Dallas Heart Study). J Am Coll Cardiol 2015;66: 1643–53.

23 D’Agostino RB Sr, Vasan RS, Pencina MJ, et al. General cardiovascular risk profile for use in primary care: the Framingham Heart Study. Circulation 2008;117: 743–53.

